# Phylogeny and Biogeography of South American Marsh Pitcher Plant Genus *Heliamphora* (Sarraceniaceae) Endemic to the Guiana Highlands

**DOI:** 10.1101/2020.04.29.068395

**Authors:** Sukuan Liu, Stacey D. Smith

## Abstract

*Heliamphora* is a genus of carnivorous pitcher plants endemic to the Guiana Highlands with fragmented distributions. We presented a well resolved, time-calibrated, and nearly comprehensive *Heliamphora* phylogeny estimated using Bayesian inference and maximum likelihood based on nuclear genes (*26S, ITS*, and *PHYC*) and secondary calibration. We used stochastic mapping to infer ancestral states of morphological characters and ecological traits. Our ancestral state estimations revealed that the pitcher drainage structures characteristic of the genus transformed from a hole to a slit in single clade, while other features (scape pubescence and hammock-like growth) have been gained and lost multiple times. Habitat was similarly labile in *Heliamphora*, with multiple transitions from the ancestral highland habitats into the lowlands. Using Mantel test, we found closely related species tend to be geographically closely distributed. Placing our phylogeny in a historical context, major clades likely emerged through both vicariance and dispersal during Miocene with more recent diversification driven by vertical displacement during the Pleistocene glacial-interglacial thermal oscillations. Despite the dynamic climatic history experienced by *Heliamphora*, the temperature changes brought by global warming pose a significant threat, particularly for those species at the highest elevations.

## 1. Introduction

The marsh pitcher plant *Heliamphora* (Sarraceniaceae) is a carnivorous plant genus endemic to the Guiana Highland across Venezuela, Guyana, and Brazil (Berry, Huber et al. 1995, Rull 2009, Rull, Huber et al. 2019). They are only found in the Pantepui region where the tepuis, or sandstone table-top mountains, rise above the surrounding lowlands (Maguire 1978, McPherson, Wistuba et al. 2011). The Pantepui region, compared to the lowland savannah, has distinct climatic, geomorphological, hydrological, and ecological features, hosting a diverse assembly of endemics with specialized adaptations (Huber 1995, McPherson 2008, Rull, Huber et al. 2019). The environments are characterized by cold temperatures, exposed surfaces, infertile substrates, and heavy precipitation (McPherson 2008, Zinck and Huber 2011).

Such harsh conditions favor diverse endemic carnivorous plant genera (e.g. *Heliamphora, Utricularia, Genlisea, Drosera* and semi-carnivorous *Brocchinia*), which thrive in the infertile areas where few non-carnivorous species are found (Brewer and Schlauer 2018, Fleischmann, Schlauer et al. 2018). Among the Pantepui endemics, the carnivorous syndrome is manifested through both morphological and molecular adaptations (Juniper, Robins et al. 1989, Matušíková, Pavlovic et al. 2018), enabling them to attract, kill and digest insect prey for additional nutrients (Brewer and Schlauer 2018). The leaves of *Heliamphora* are not only photosynthetic but also highly modified pitchers with structures specialized in carnivorous functions (McPherson, Wistuba et al. 2011). *Heliamphora* produces nectar as attractant in the nectar spoon above the pitcher opening (Płachno and Muravnik 2018), enticing prey to fall into the pitcher filled with digestive liquid. The prey then drown and are digested, allowing the plant to absorb the nutrients rich in nitrogen and phosphorous (Maguire 1978, Juniper, Robins et al. 1989). Unlike other pitcher plants (*Sarracenia, Darlingtonia, Nepenthes* and *Cephalotus*) that have lids or hoods covering the pitcher, *Heliamphora* do not have similar structures to prevent rainwater from entering the adult pitchers. Instead, the adult pitchers of *Heliamphora* are adapted to the high rainfall in the Pantepui by allowing excess rainwater to flow through a drainage hole or slit below the pitcher opening (McPherson, Wistuba et al. 2011). Additionally, the retentive hairs above the digestive fluid and around drainage structures act as filters that prevent captured prey from escaping from the pitchers (Jaffe, Michelangeli et al. 1992).

*Heliamphora* populations are generally confined to the montane or tepui summits at elevations above 1500m (McPherson, Wistuba et al. 2011). While some species are distributed across a group of near-by tepuis (massifs or tepui chains), more than half of the genus only occurs in single tepuis, valleys, canyons, or upland swamps (McPherson, Wistuba et al. 2011). Poor seed dispersibility and presence of lowland savannah acting as dispersal barrier may have contributed to this high degree of endemism. It had been suggested that *Heliamphora* might have diversified through allopatric speciation as the once continuous highland habitat gradually fragmented over time through erosion (Maguire 1978, Huber 1987, McPherson, Wistuba et al. 2011, Huber 2018).

In this study, we estimated the time-calibrated phylogeny of *Heliamphora* in order to resolve relationships within the genus and trace its diversification over space and time. We tested the relationship between geographic and genetic distance, with the expectation that low dispersal ability would result in close relatives living in close proximity. We also estimated the history of changes in habitats using ancestral state reconstruction to test for directionality in transitions (i.e. highland vs. lowland). Finally, we reconstructed key morphological characters to identify traits diagnostic of clades. In interpreting these results, we considered possible modes of speciation and discussed the possible impacts of global climate change on the future of *Heliamphora*.

## 2. Materials and methods

### 2.1 Taxon Sampling

Recent treatments of *Heliamphora* recognize 23 species, most of which were described since 2000 (McPherson, Wistuba et al. 2011). We sampled 22 described *Heliamphora* species (all except *H. macdonaldae*) plus 2 undescribed taxa from collectors or commercial sources for the phylogenetic analysis (Appx. A Sample Accession Numbers and Sources). We confirmed species determinations by comparing the morphological traits of these living collections to the original descriptions (Bentham 1840, Nerz and Wistuba 2000, Wistuba, Harbarth et al. 2001, Wistuba, Carow et al. 2002, Carow, Wistuba et al. 2005, Wistuba, Carow et al. 2005, Fleischmann, Wistuba et al. 2009). We also sampled five taxa (*Sarracenia leucophylla, S. pittacina, Darlingtonia californica, Roridula gorgonias* and *Actinidia arguta*) to serve as outgroups (Appx. A Sample Accession Numbers and Sources). Previous work had supported the monophyly of Sarraceniaceae (comprising the three genera, *Sarracenia, Heliamphora* and the monotypic *Darlingtonia*) and shown that Roridulaceae+Actinidiaceae were sister to the family (Ellison, Butler et al. 2012, Magallón, Gomez-Acevedo et al. 2015).

### 2.2 DNA Extraction, Amplification and Sequencing

All *Heliamphora, Sarracenia psittacina*, and *Darlingtonia californica* were grown under controlled environmental conditions (photoperiod 15hr per day; temperature range 12-24 °C; relative humidity >80%) inside of growth chambers under artificial lights with temperature/humidity regulating units in the University of Colorado Boulder before DNA extraction. DNA was extracted from 0.5-1.0 grams of fresh, young, and unopened pitchers using the CTAB protocol (Doyle and Doyle 1987).

We amplified and sequenced three nuclear genes (ITS, 26S, PHYC) using polymerase chain reaction (PCR) amplification and Sanger Sequencing. The markers had shown potential in resolving interspecies relationships in Sarraceniaceae from a previous study (Ellison, Butler et al. 2012). The ITS region was amplified using the primers ITS4 (White, Bruns et al. 1990) and ITS5 (Ristaino, Madritch et al. 1997) or ITSLEU (Baum, Small et al. 1998). The 26S region was amplified using overlapping primer sets N-nc26S1/1229rev, N-nc26S4/1499rev, N-nc26S4/2134rev, nc26S9/3058rev, and nc26S9/3331rev (Kuzoff, Sweere et al. 1998). PHYC was amplified using primers cdo (Mathews and Donoghue 1999) and int1F (Wurdack and Davis 2009). The PCR conditions and recipes are provided in Appendix B (PCR Mix Recipes and Conditions). Sequences for three of the outgroups (*S. leucophylla, R. gorgonias*, and *A. arguta*) were available from previous studies (Ellison, Butler et al. 2012, Lo□fstrand and Scho□nenberger 2015) and were downloaded from GenBank (Appx. A).

### 2.3 Phylogenetic Analysis

Nucleotide sequences were *de novo* assembled from chromatogram files in Geneious 11.1.5 (https://www.geneious.com). All sequences were first aligned under default settings (Geneious global alignment with free end gaps and 93% similarity cost matrix). Uncertain positions, where peaks corresponding to different bases were similar in height, were scored as ambiguous (e.g., W for A or T). The alignment was trimmed at the two ends so that all sequences were mostly overlapping and similar in length. Bayesian phylogenetic analysis was carried out in BEAST 2.5.2 (Bouckaert, Heled et al. 2014). Nucleotide substitution models were chosen for each marker by running alignments in jModeltest 2.1.10 with the Bayesian information criterion (Guindon and Gascuel 2003, Darriba, Taboada et al. 2012). GTR + G + I, TN93, and HKY models were selected for 26S, ITS and PHYC, respectively. We used a birth-death process for diversification priors. We constrained *R. gorgonias* and *A. arguta* to form a clade given the lack of complete genetic data in this study and the well-supported relationships from previous studies (Ellison, Butler et al. 2012, Lo□fstrand and Scho□nenberger 2015). A chain length of 30 million generations sampling every 1000 trees was used to estimate the model parameters. We verified that all parameters had ESS values above 200 using Tracer 1.7.1 (Rambaut, Drummond et al. 2018). TreeAnnotator 2.4.7 was used to generate a maximum clade credibility (MCC) tree with posterior probabilities (Bouckaert, Heled et al. 2014). The MCC tree was first visualized in FigTree 1.4.4 (http://tree.bio.ed.ac.uk/software/figtree/), then further edited in Affinity Designer 1.7.1. We also estimated individual gene trees in RAxML 8.2.10 (Stamatakis 2014) using the GTRGAMMA model with 1000 bootstraps using CIPRES Portal (Miller, Pfeiffer et al. 2010).

### 2.4 Divergence Time Calibration

Two secondary calibration points were used to estimate node ages in Sarraceniaceae, further supported by several fossils dated prior to those estimations. A fossil named *Archaeamphora longicervia* Hongqi Li was first assigned to Sarraceniaceae due to its structural similarity to *Heliamphora* and *Sarracenia purpurea* (Li 2005). However, its early Cretaceous age (~125Ma) made it unlikely to be an early representative of Sarraceniaceae since several studies with robust fossil calibrations found the family to be younger than 90 Ma (Magallón, Gómez-Acevedo et al. 2015, Tank, Eastman et al. 2015, Wong, Dilcher et al. 2015, Rosea, Kleistb et al. 2018). Several fossils had been described in Actinidiaceae, with *Parasaurauia allonensis* (Keller, Herendeen et al. 1996), *Paradinandra suecica* (Schönenberger and Friis 2001), and *Glandulocalyx upatoiensis* (Schönenberger, Balthazar et al. 2012) dating back in the late Santonian (~84 Ma). In Roridulaceae, well preserved *Roridula* carnivorous leave fossils were discovered in Eocene Baltic amber dating to less than 47 Ma, indicating that Roridulaceae emerged before 47Ma but probably after 84Ma (Sadowski, Seyfullah et al. 2015). The robust, fossil calibrated angiosperm phylogeny from Magallón et al. (2015) estimated the mean ages of Sarraceniaceae+Roridulaceae/Actinidiaceae and Roridulaceae+Actinidiaceae to be 88.09 and 83.58 Ma, respectively (Magallón, Gómez-Acevedo et al. 2015). Given the concordance with the fossil dates, we chose to use these estimates from the angiosperm-wide analysis (Magallón, Gómez-Acevedo et al. 2015) as secondary calibrations. We entered 88.1 (95%HPD:85.2-91.0) and 83.6 (95%HPD:82.8-86.8) Ma as priors for those nodes indicated above.

### 2.5 Mantel Test for Phylogenetic Signal in Geographic Distribution

Methods centered on biogeographic models, such as dispersal-extinction-cladogenesis (DEC) (Ree, Moore et al. 2005, Ree and Smith 2008) and its extensions (Matzke 2013), have often been used to infer range shifts (Matzke 2014, Klaus, Morley et al. 2016, Fuchs, Pons et al. 2017) and and could be applied to *Heliamphora*. However, conceptual and statistical issues were recently identified that make likelihood comparisons between DEC and DEC+j models invalid (Ree and Sanmartin 2018). Although alternatives, like ClaSSE (Goldberg and Igic 2012) corrected the issue by modeling cladogenesis as time-dependent probabilities, they are too parameter rich to be applied to *Heliamphora*, given its small size and complex local distribution patterns. Instead, we implemented a simpler Mantel test in R 3.6.1 to test for correlation between genetic and geographical distance between each species (Diniz-Filho, Soares et al. 2013). We predicted that the potential limited dispersal ability of *Heliamphora* (Cross, Davis et al. 2018) would lead to a strong phylogenetic signal in geography, with close relatives tending to occur in close proximity. We used the estimated phylogeny to produce the genetic distance matrix. We then used centroid coordinates (Appx. C Species Distributions) from each species’ known distribution in McPherson et al. (2011) to produce the geographical distance matrix.

### 2.6 Ancestral State Estimation

A range of morphological features, such as aspects of pitcher size, form and pubescence, are diagnostic for *Heliamphora* species (McPherson, Wistuba et al. 2011). We estimated the evolutionary histories of three such morphological characters: drainage hole, scape pubescence, and growth form. As some of these traits are absent in the outgroups, we first pruned the phylogeny to include only *Heliamphora*. Trait evolution was estimated using stochastic mapping with the function “make.simmap” in phytools 0.6-99 (Revell 2012). All characters except for scape pubescence were scored as binary based on species descriptions (Bentham 1840, Steyermark 1951, McPherson, Wistuba et al. 2011). Scape pubescence was scored as “glabrous”, “pubescent” and “glabrous+pubescent” as some species are found to have both pubescent and glabrous scapes. After stochastic mapping, we combined later two states as “pubescent” as we disregarded transitions between “pubescent” and “glabrous+pubescent”. For every character, we compared for two transition rate models: the equal rates (ER) model and the all rates different (ARD) model (Pagel 1994, Schluter, Price et al. 1997). The best fit model for each character was selected based on the lowest AIC scores and used for stochastic mapping. We also examined habitat shifts in *Heliamphora* using the same approach. We scored habitat types as either Upper or Lower Pantepui (highland vs. lowland habitats) based on a taxon’s lowest distribution. We used 1500m as our threshold elevation for habitat types because drastic shifts in local climate, geomorphology, and community composition were observed at this elevation (Huber 1987, Rull, Huber et al. 2019).

## 3. Results

### 3.1 Phylogenetic Analysis & Divergence Time Estimation

Our analysis supported the sister clade relationship between *Sarracenia* and *Heliamphora*, in accordance with previous studies (Neyland and Merchant 2006, Ellison, Butler et al. 2012, Stephens, Rogers et al. 2015). Within *Heliamphora*, we recovered well supported subclades found in the western (W) and the eastern (E) Pantepui region. For the purpose of discussion, we divided the species-rich eastern clades into four (E1, E2a, E2b, E3) (Fig. 1). In *Heliamphora*, the monophyly of all sub-clades and even most of the interspecies relationships were well supported (posterior probability >0.99) by our combined nuclear data set. Relationships within *Heliamphora* showed strong geographic structure. For example, the W clade was restricted to the western Pantepui and the E3 clade to the Eastern Tepui Chain (Fig. 1). Both 26S (3251bp) and ITS regions (676bp) supported the monophyly (bootstrap scores BS > 80) of the W, E1, E2 and E3 clades (Appx. D Individual Gene Trees Estimated Using Maximum Likelihood). PHYC (787bp) provided good support (BS > 85) for the monophyly of the sub-clades E2a and E2b (Appx. D). Using the two previously described node calibrations, we estimated that *Sarracenia* and *Heliamphora* split ca. 14 mya, and that the various subclades diverged between 6 and 2 mya (Fig. 1).

**Figure 1.**
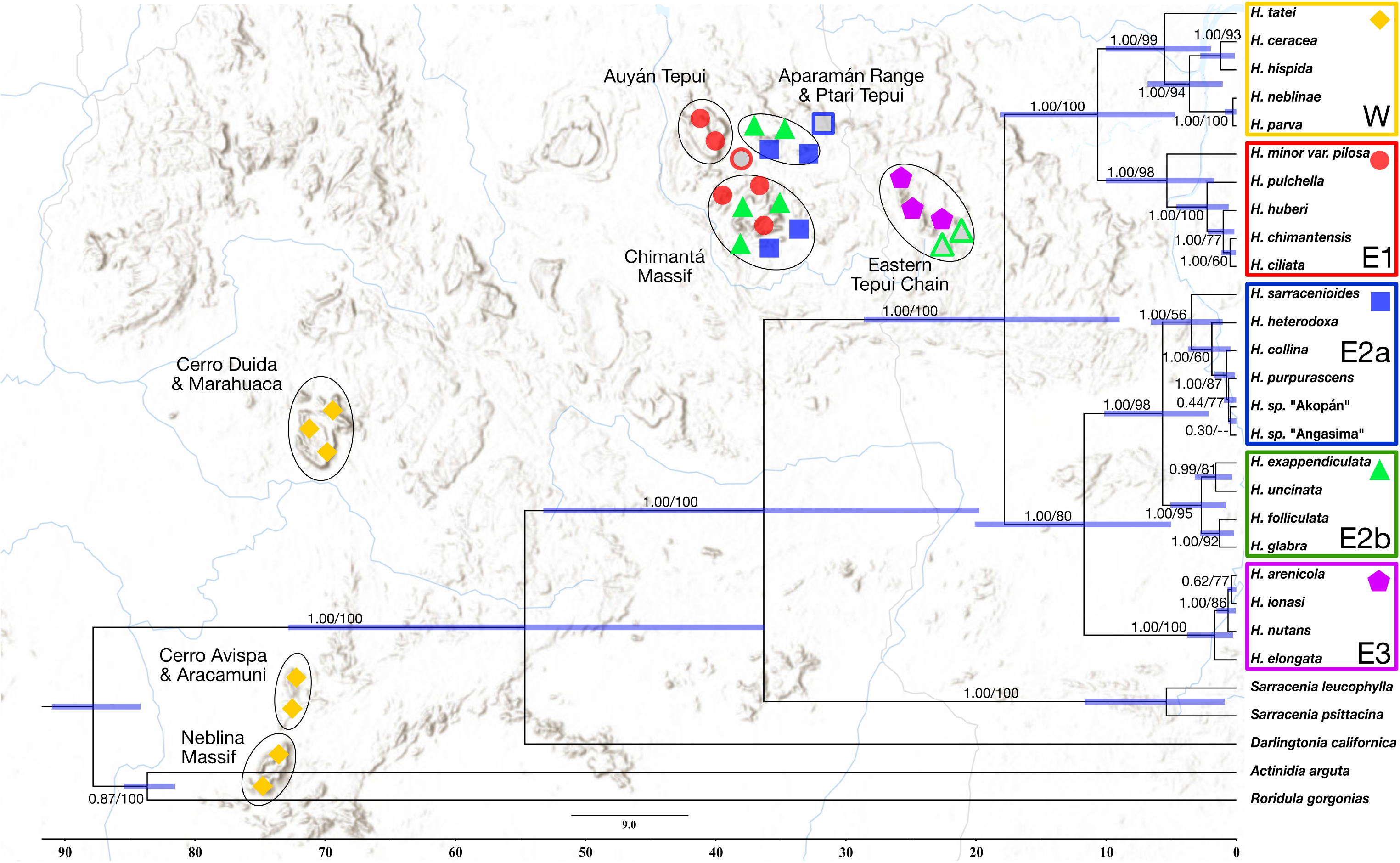
Phylogeny and Distribution of *Heliamphora*. The phylogeny was based on combined nuclear sequences (ITS, 26S, and PHYC) with Bayesian posterior probability and ML bootstrap score indicated at the branches of each major clade. Major clades were marked with colors and shapes both shown on the phylogeny and distribution. Highland species were shown with solid shapes while lowland species were shown with open shapes.

### 3.2 Mantel Test and Biogeography

Our null hypothesis (no correlation between genetic and geographic distance) was rejected by the Mantel test (p<0.001, r=0.43), suggesting that genetic distances are correlated with geographic distances. This result indicates that close relatives tend to be distributed closely and distantly related species farther apart. The geographic structuring is apparent from the the phylogeny, where species in each subclade were clustered within the Pantepui (Fig. 1). One exception was the E2b clade, which is widespread in the eastern region. Also, the western clade (W) is monophyletic but nested within the genus, making eastern clade (E1, E2, and E3) paraphyletic.

### 3.3 Ancestral State Estimation

Based on the AIC scores, the ER model was selected to estimate the ancestral states of the drainage structure and scape pubescence (Fig. 2). The ARD model was selected for the evolution of growth form and habitats. For the drainage structure, we estimated a single transition from drainage hole to slit [median: 1 with 95% credibility interval (CI): 0-2 and likely no reverse transitions] (Fig. 2). The median transition numbers from glabrous to pubescent scape and vice versa were estimated to be 2 (CI: 0-6) and 2 (CI: 1-4), respectively (original 3 state transition ASR see Appx. E). The adaptation and loss to hammock-like clumpy growth form was inferred to be frequently gained and lost with median numbers of transitions estimated to be 36 (CI: 14-80) and 37.5 (CI: 14-80), respectively (Appx. E Ancestral State Reconstruction for Growth Forms and Habitats). Similarly, the transitions between highland and lowland habitat was inferred to be frequent from both directions [34 (CI: 13-74) and 34 (CI: 12-77)] (Appx. E). We suspect that these extremely high numbers of inferred transitions reflects high rates estimated along short branches in the tree (see Discussion 4.1.2).

**Figure 2.**
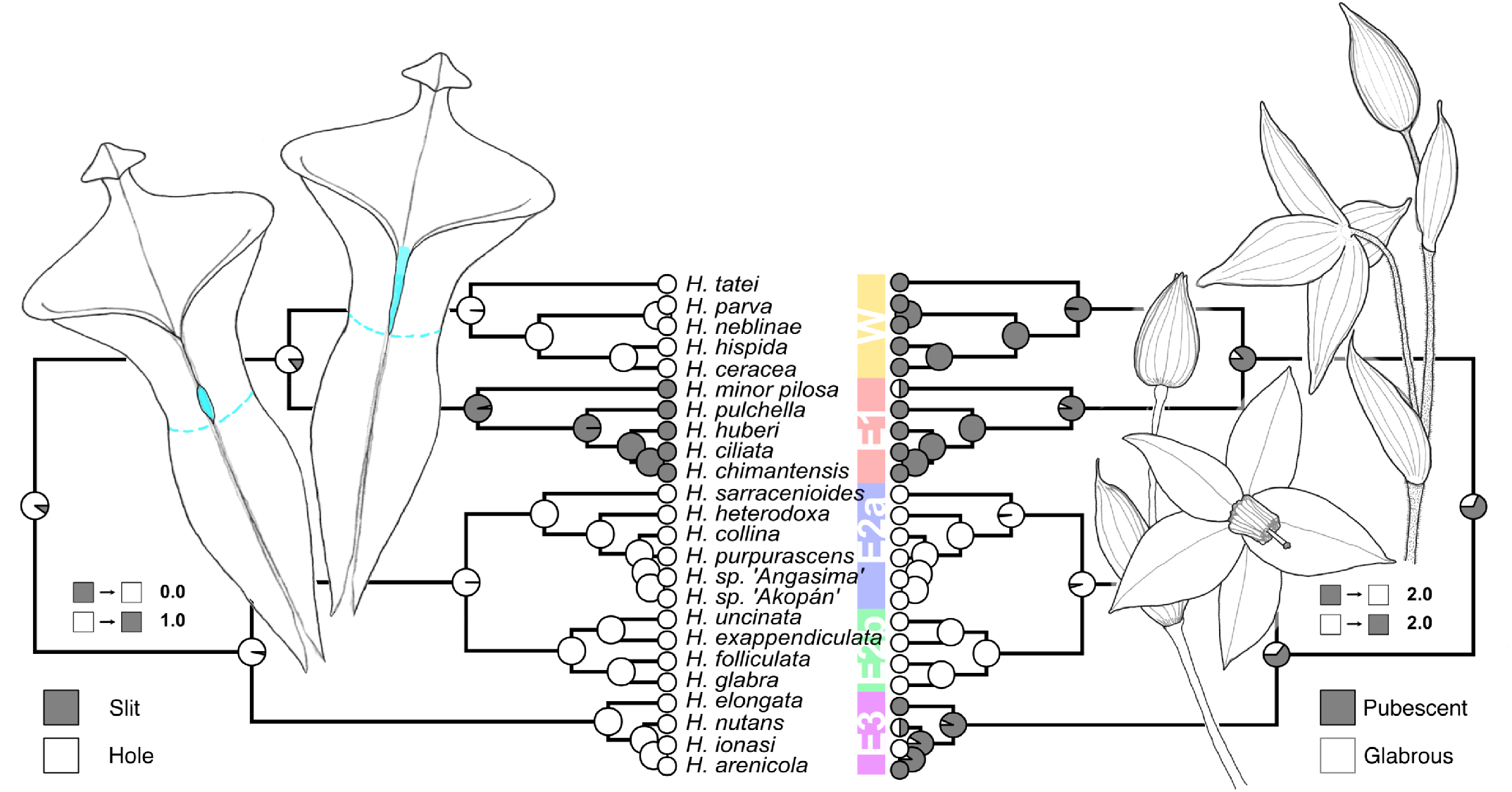
Ancestral state reconstruction of drainage structures (left) and scape pubescence (right) using stochastic character mapping. Median numbers of transitions between states were shown in the legends. In the simplified *Heliamphora* adult pitchers diagram modified from McPherson, Wistuba et al. (2011), drainage hole, drainage slit were filled with cyan color, and digestive fluid levels were marked with cyan dotted lines. Diagram of *Heliamphora* inflorescences with pubescent and glabrous scapes were adapted from Fleischmann, Wistuba et al. (2009). Scapes of *H. nutans* and *H. minor* are either pubescent or glabrous, represented with half circles.

## 4. Discussion

The previous family-level analysis (Ellison, Butler et al. 2012) recovered a monophyletic *Heliamphora* (with six taxa sampled), but with low levels of support for relationships within the genus. With newly obtained sequence data for all *Heliamphora* but *H. macdonaldae*, we were able to resolve relationships among most of the taxa with strong support. Using updated secondary calibration points, we recovered age estimates that were generally older than those in previous studies (Ellison, Butler et al. 2012, Rose, Kleist et al. 2018) but in accordance with known fossil records for Ericales (Sadowski, Seyfullah et al. 2015). In the context of this new phylogeny, we discuss patterns of trait evolution and possible biogeographical histories that might have shaped present-day *Heliamphora* distributions along with the implications of global warming for the future of this clade.

### 4.1 Evolution of Morphological and Ecological Traits

#### 4.1.1 Drainage Structures & Scape Pubescence

Drainage structures, in the form of holes or slits in the pitchers, allow *Heliamphora* to regulate fluid levels by draining excess rainwater out while retaining captured prey (McPherson, Wistuba et al. 2011). No homologous drainage structures are present in *Sarracenia* or *Darlingtonia*, suggesting that these structures evolved in ancestral *Heliamphora* populations, presumably as an adaptation to the perhumid climate in the Pantepui region. The drainage slit (Fig. 2) is found exclusively in the E1 clade where a narrow V-shaped slit extends from pitcher opening to the upper half of the ventral suture. In the rest of *Heliamphora*, the drainage hole is present and generally positioned at the lower half of the ventral suture. Our analysis revealed that the hole is the ancestral state in *Heliamphora* and was further modified into drainage slit (Fig. 2).

Scape pubescence was another character of taxonomic importance, although our analysis indicated it has a complex history. We inferred that a pubescent scape is ancestral in *Heliamphora* but was further lost in the ancestral population leading to E2 clade (Fig. 2). Moreover, our analysis suggested that pubescent scape might have been independently lost in *H. minor*, *H. nutans*, and *H. ionasi*, showing this trait could be frequently lost during diversification.

#### 4.1.2 Highland to Lowland Transitions

Most of *Heliamphora* species occur exclusively in the upper Pantepui region at elevations higher than 1500m (Fig. 2). However, a few species could be found in the lower elevations below 1500m (Appx. C). *H. glabra* and *H. heterodoxa* are distributed from upper Pantepui into the lower elevations at around 1200m while *H. ciliata* is the only species that occur exclusively in the lowland at 900m. Our analysis suggested that the most likely ancestral state across the clade is highland (Appx. E), but we also estimated frequent transitions between upper and lower Pantepui habitats (see Results 3.3). Given that only three taxa occur in lower elevations, we consider the high number of inferred transitions to be a spurious result relating to the short branch lengths separating sister taxa occurring in different habitats (e.g. *H. ciliata* in Lower Pantepui vs. *H. chimantensis* in Upper Pantepui). The presence of such contrasts separated by short time periods results in high estimated transition rates in Markov models of trait evolution (Pagel, Meade et al. 2004).

#### 4.1.3 Hammock-like Clump Growth Habit

Diverse growth habits are found in *Heliamphora*, possibly as adaptations to the local environments (McPherson, Wistuba et al. 2011). Some species occurring in exposed summit environment tend to exist in extensive clonal populations, forming hammock-like clumps up to several meters in diameter (Wistuba, Carow et al. 2002, Wistuba, Carow et al. 2005) while other species growing in densely vegetated montane habitat tend to grow singly or even stem forming (McPherson, Wistuba et al. 2011). Our ancestral state reconstruction suggested that growth form was not highly conserved, with multiple shifts along the phylogeny (Appx. E). Nevertheless, ancestral states were uncertain throughout, leaving it unclear when these transitions have occurred. Future studies would be valuable to test the ecological drivers of this growth habit variation.

### 4.2 Evolutionary and Biogeographical Histories

After the ancestral lineage split into two major lineages giving rise to W/E1 and E2/E3 in the early Miocene, *Heliamphora* underwent three waves of diversification (Fig. 3). The first event occurred during Late Miocene when E2 and E3 diversified shortly followed by the splitting of W and E1. During the transition between Miocene and Pliocene, the second wave took place when E2 further split into E2a and E2b, followed by further diversification of W and E1 clades. The split of *H. tatei* (together with its potential sister *H. macdonaldae*, not sampled here) from the rest of W clade might have taken place due to lack of connectivity between Duida–Marahuaca Massif and Neblina–Aracamuni Massif. Similarly, the divergence of *H. minor* from the rest of E1 clade could have resulted from allopatric speciation when Auyán Tepui and Chimantá Massif lost their connectivity (Rull and Nogué 2007). The last wave of diversification occurred from the late Pliocene into Quaternary when rapid speciation took place within all *Heliamphora* clades.

**Figure 3.**
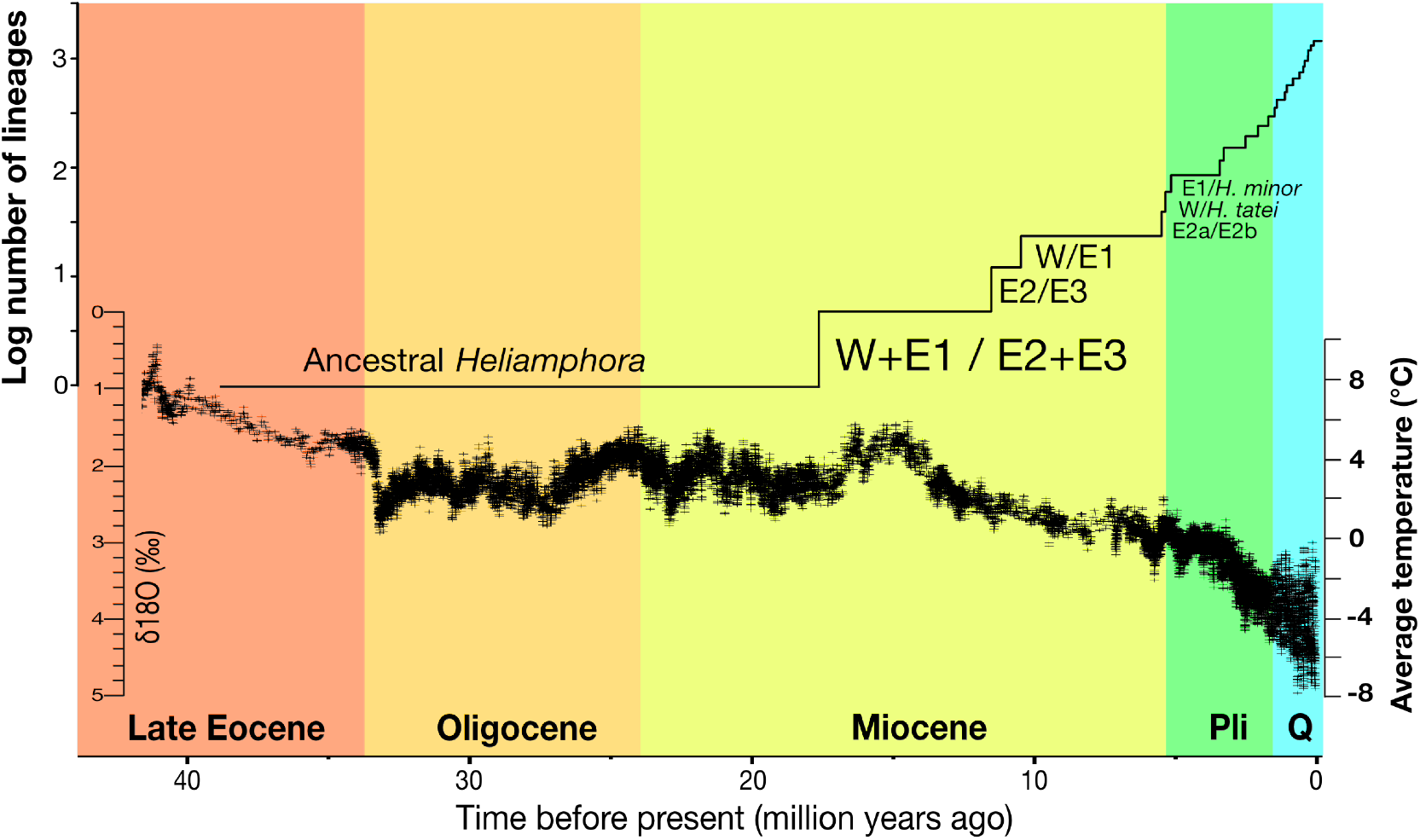
Lineage through time plot on log-scale shows the diversification trend of *Heliamphora* in relation to the Climate Trend in the Past ~45 Ma. Major clades in *Heliamphora* might have emerged by Pliocene (Pli.) when global average temperature drastically decreased and more suitable habitat became available in the Guiana Highland region. During the Pleistocene, *Heliamphora* diversified rapidly, possibly facilitated by glacial-interglacial oscillations. The climate curve data was obtained from Zachos et al. (2001) and Zachos et al. (2008) based on deep-sea benthic foraminiferal oxygen-isotope curve (δl80‰).

The high degree of endemism in *Heliamphora* clades suggests that vicariance or short distance dispersal could have played an important role in shaping the present-day diversity. As supported by our Mantel test, closely related species showed strong geographic clustering. The distributional pattern was consistent with the hypothesis proposed by Maguire (1970) that many Pantepui endemics, like *Heliamphora*, could have once occurred on a more continuous highland habitat but later diversified as the highland gradually became eroded, fragmented and eventually separated (Maguire 1970). Still, the nested position of the west (W) clade within the eastern clades points to a more complex biogeography history, potentially involving rare long distance dispersal from the eastern region.

While biogeographic events such as vicariance and dispersal may account for the origins of major clades, more recent climatic oscillations could have contributed to recent diversification events within the genus. Available paleoclimatic data (Zachos, Dickens et al. 2008, Hansen, Sato et al. 2013) suggest that the Pantepui region might have experienced a warmer climate during the early Miocene but entered rapid cooling phase since mid-Pliocene (Fig. 3). By the Pleistocene, thermal oscillations initiated and average temperature shifts of more than 8 °C frequently occurred during the transition between glacial and interglacial periods (Rull and Vegas-Vilarrúbia 2019). Although the tepui summits were never glaciated during glacial periods (Rull, Montoya et al. 2019), the drastic temperature shifts would have profound effect on the distributions of *Heliamphora* in the Pantepui (Steyermark and Dunsterville 1980). During the glacial periods, the highland habitats could have shifted downward (Rull and Vegas-Vilarrúbia 2019), allowing *Heliamphora* populations to descend, colonize surrounding lower elevations and open new migrational routes between tepui summits (Steyermark and Dunsterville 1980, Rull 2005, Rull and Nogué 2007). In the warmer interglacial periods, *Heliamphora* populations may have migrated to higher elevations where suitable habitats had advanced back up (Rull 2005, Rull and Nogué 2007). Pollen analysis of peat outcrops deposited on tepui summits during the warmer, more stable Holocene revealed *Heliamphora* and other Pantepui endemic communities had shifted distributions for at least a few hundred meters in response to temperature shifts (Rull 2004, Rull, Montoya et al. 2011). This indicated that more intense vertical displacements would have potentially occurred during the stronger thermal oscillations in Pleistocene, allowing *Heliamphora* to descend downward to up to 1,100m into the lowlands (Farrera, Harrison et al. 1999, Rull and Nogué 2007, Rull and Vegas-Vilarrúbia 2019).

### 4.3 Evolutionary Future in a Warming Climate

Thus far, *Heliamphora* has largely remained unthreatened thanks to the remoteness and inaccessibility of their native habitats with few historical anthropogenic disturbances (McPherson, Wistuba et al. 2011, Safont, Rull et al. 2016, Bevilacqua, Señaris et al. 2019). However, ongoing effect of global warming poses a severe threat to *Heliamphora* (Nogué, Rull et al. 2009, Rull, Nogué et al. 2019). The global temperature is projected to increase 2-4 °C by the end of the century (JT, Y et al. 2001), rendering it hotter than any interglacial periods ever experienced during the Pleistocene (Fig. 3). Such projected temperature increase would cause *Heliamphora* suitable habitats to shift upward 300-600m in elevation (Rull, Nogué et al. 2019). *Heliamphora* species restricted to tepui summits with smaller areas and lower elevations (e.g. *H*. sarracenioides, *H. purpurascens* and *H. spec. nov. ‘Angasima’*) would lose suitable habitats with no higher elevation to migrate to, and eventually go extinct (Appx. C). Other *Heliamphora* species, if they survive the warming by rapidly dispersing and colonizing upward, would still suffer severe habitat loss by 25-98% due to reduction of available surface area towards higher elevations (Nogué, Rull et al. 2009).

## Supporting information

Appx A

Appx B

Appx C

Appx D

Appx E

