## Supplementary figures and images for "Phylogeny and Biogeography of South American Marsh Pitcher Plant Genus *Heliamphora* (Sarraceniaceae) Endemic to the Guiana Highlands"

### Appx D

#### Appendix C.

#### Individual Gene Phylogeny Estimated by Maximum Likelihood

a. 26S

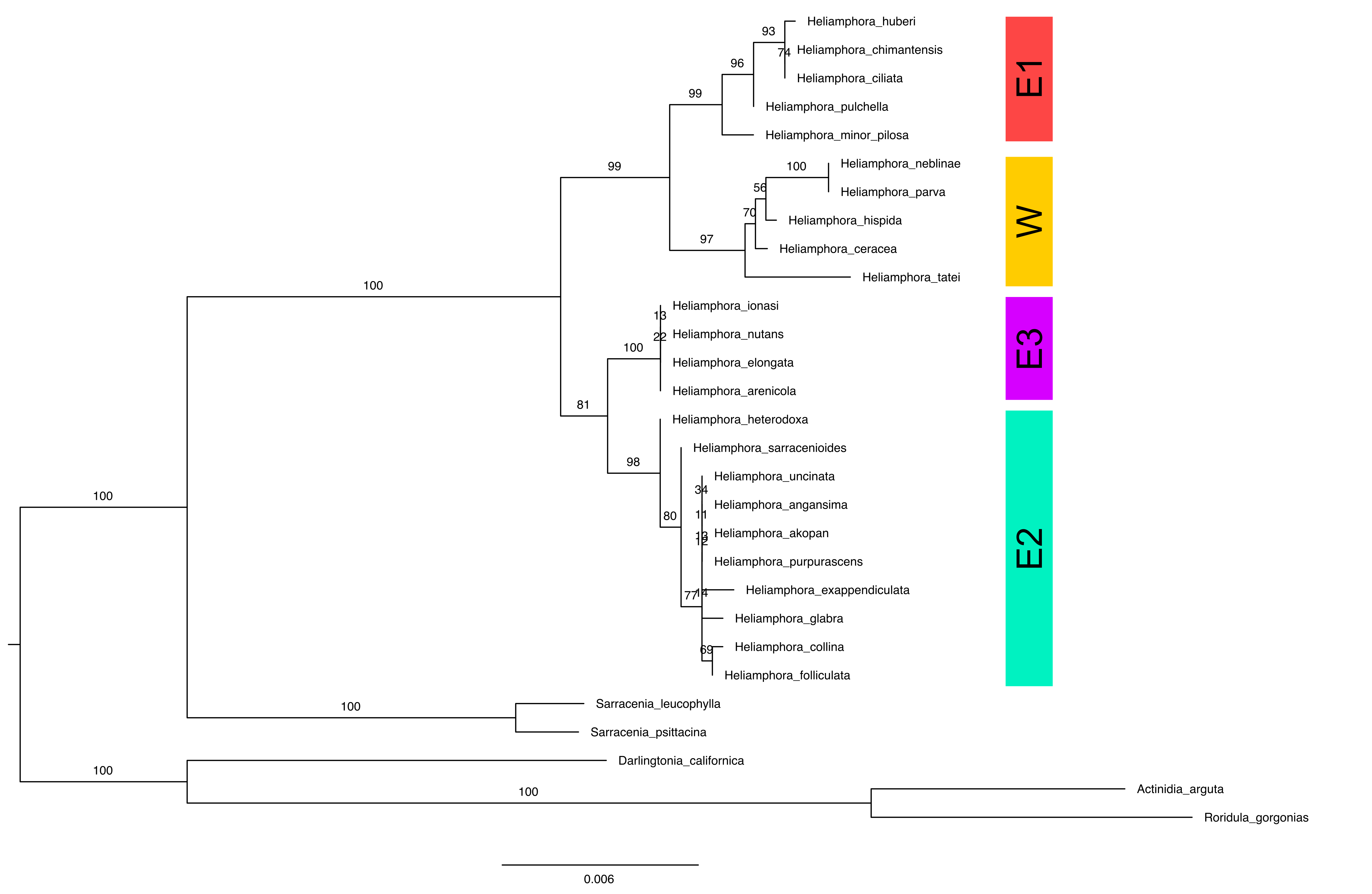

a. ITS

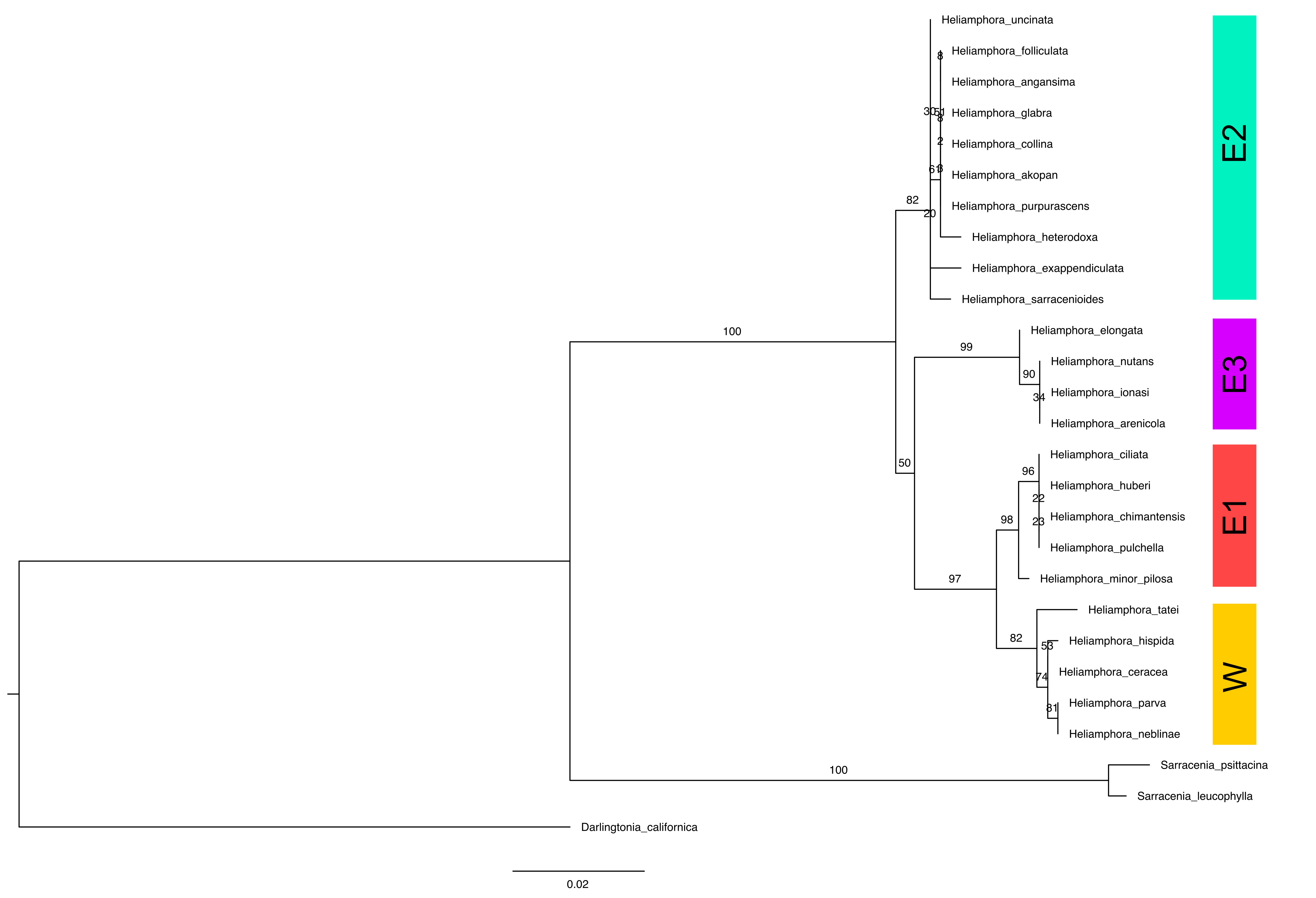

b. PHYC

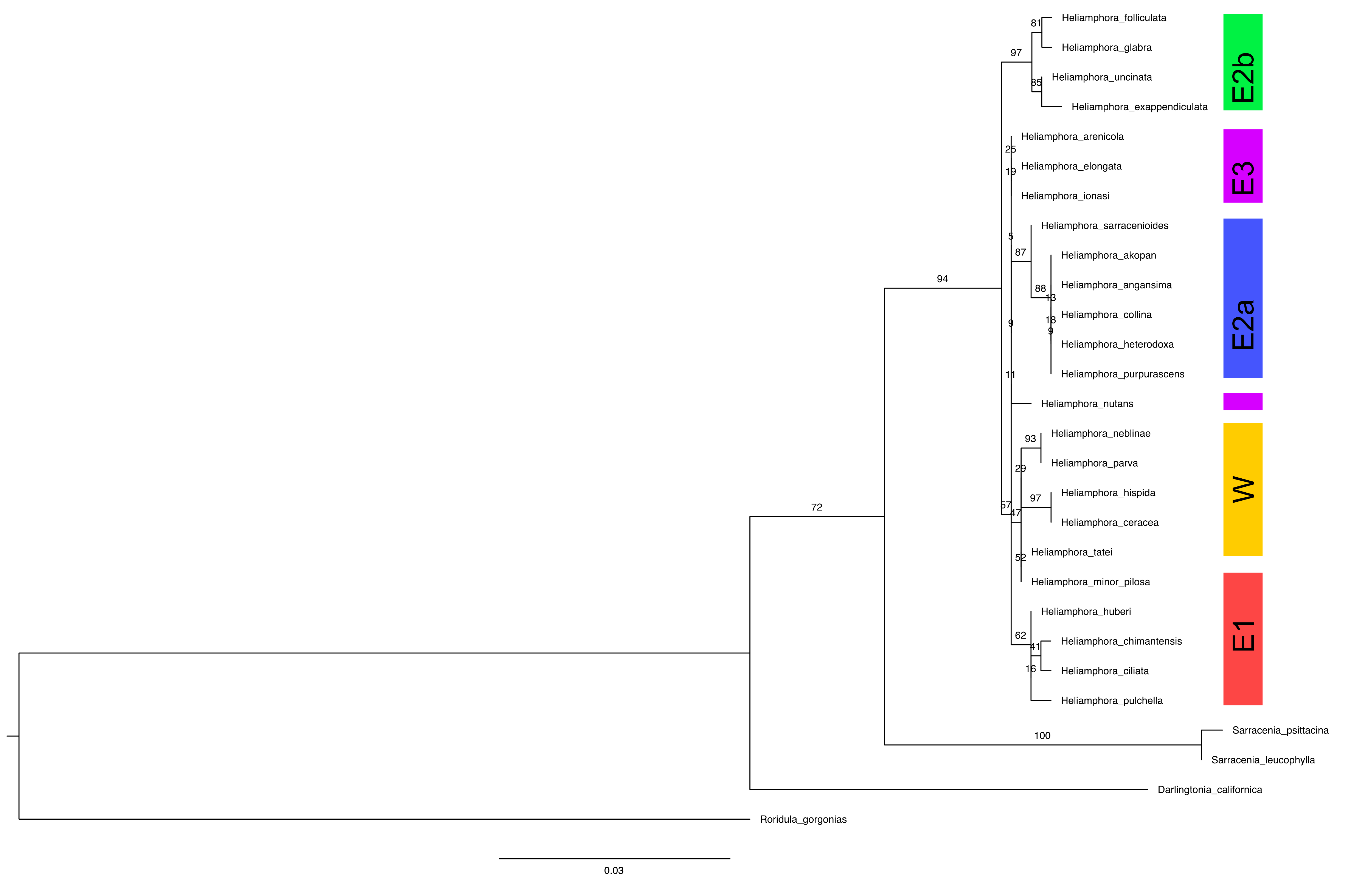
