## Supplementary material for "Phylogeny and Biogeography of South American Marsh Pitcher Plant Genus *Heliamphora* (Sarraceniaceae) Endemic to the Guiana Highlands": Appx E

### Appendix E.

### Ancestral State Reconstruction of Growth Form and Habitats

a. Hammock-like Clumpy Growth Form (Clumpy vs. Not Clumpy)

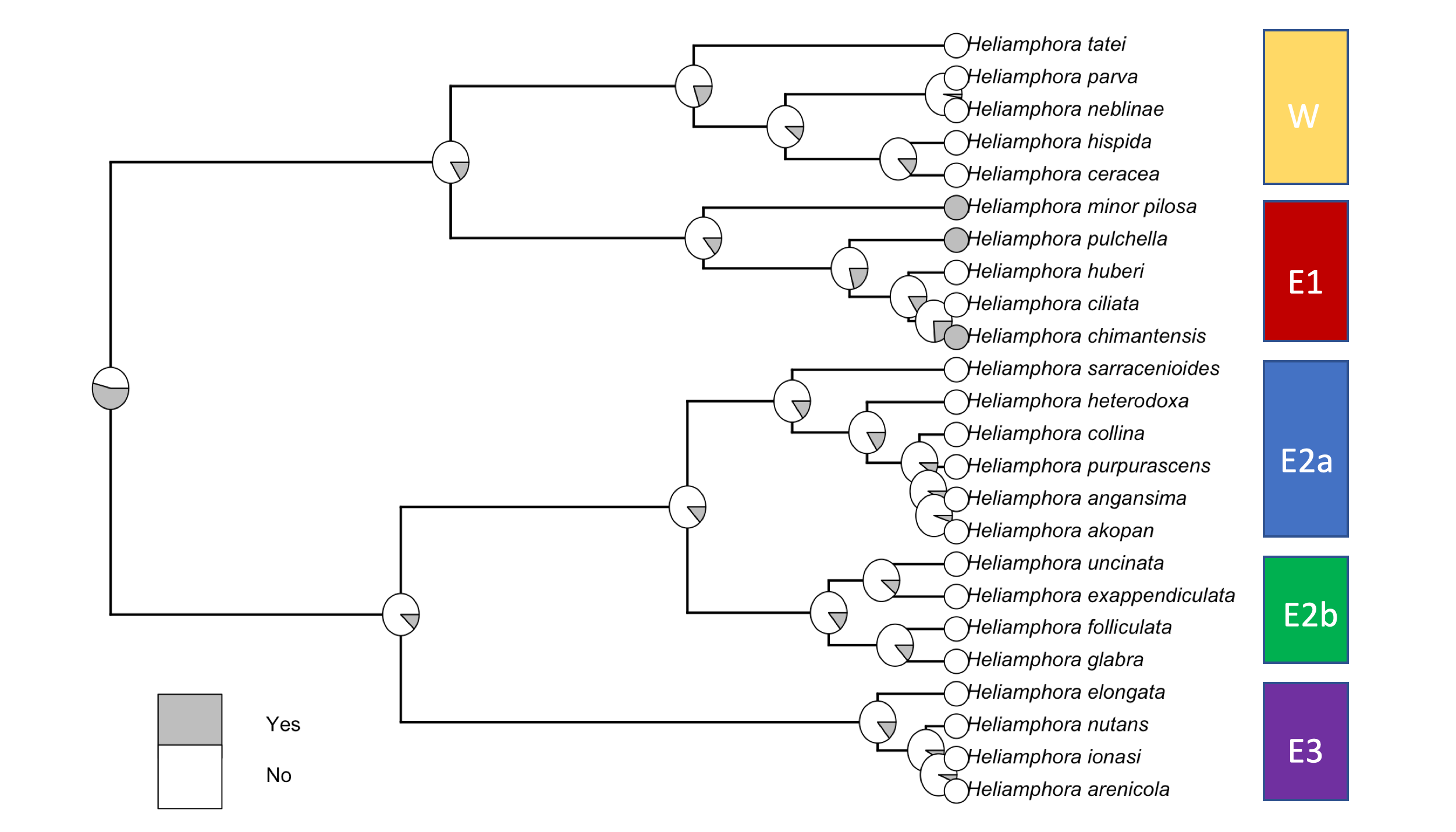

b. Habitats (Upper vs. Lower Pantepui)
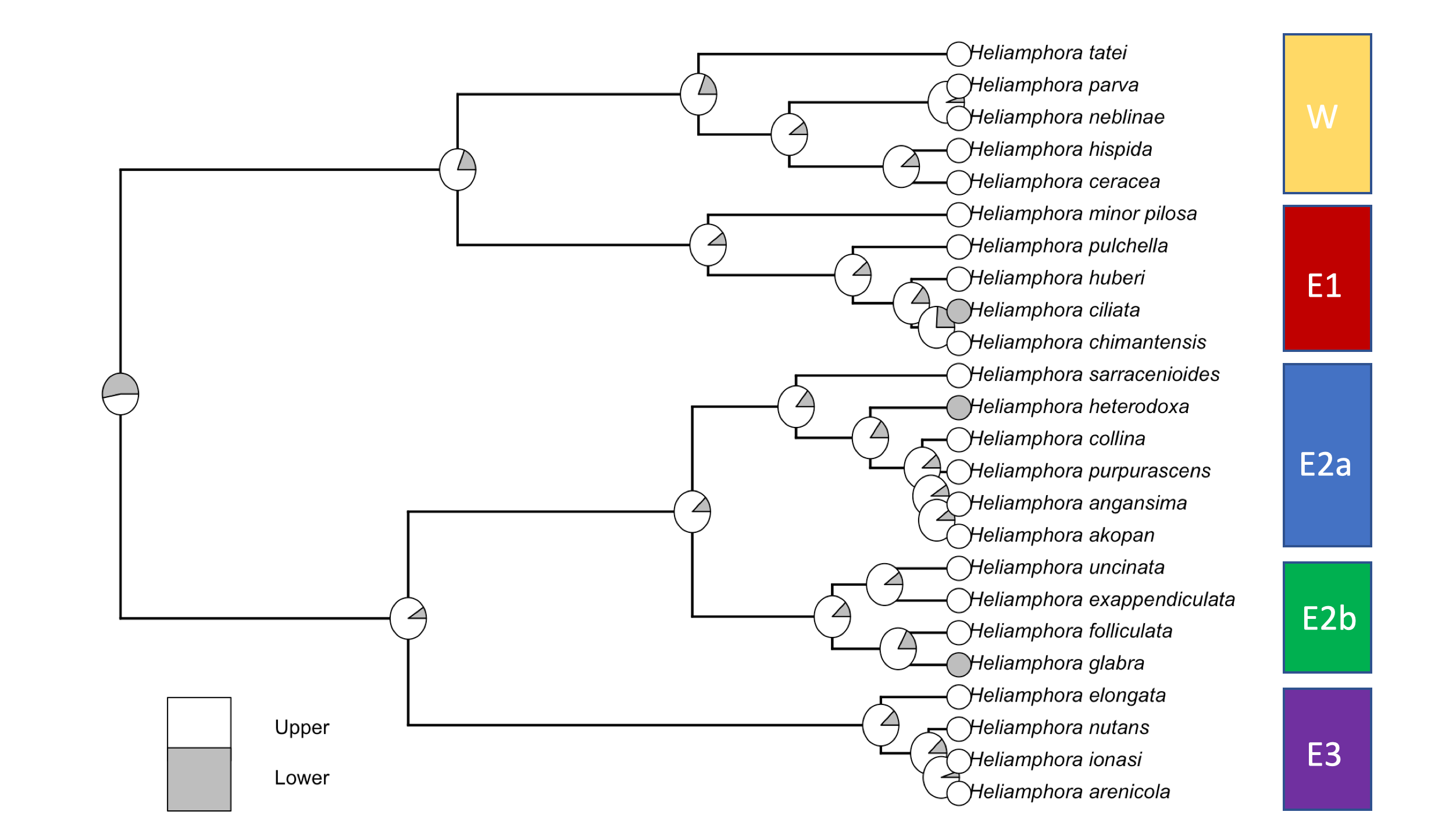

c. Scape Pubescence (Glabrous, Pubescent, Pubescent+Glabrous)

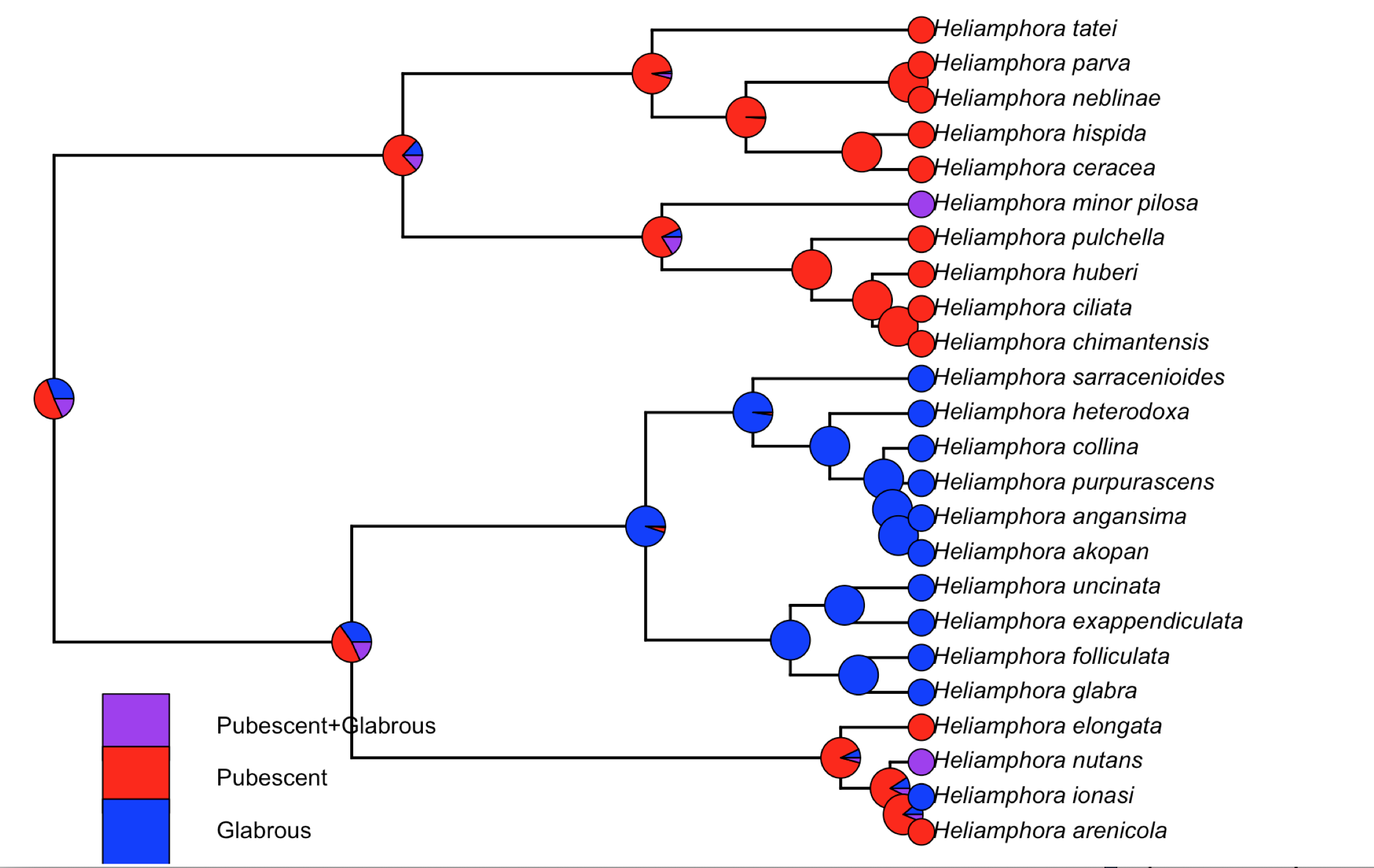
